# Ratatosk – Hybrid error correction of long reads enables accurate variant calling and assembly

**DOI:** 10.1101/2020.07.15.204925

**Authors:** Guillaume Holley, Doruk Beyter, Helga Ingimundardottir, Snædis Kristmundsdottir, Hannes P. Eggertsson, Bjarni V. Halldorsson

**Affiliations:** deCODE genetics/Amgen Inc., Reykjavík, Iceland; School of Technology, Reykjavik University, Reykjavík, Iceland

## Abstract

**Motivation:** Long Read Sequencing (LRS) technologies are becoming essential to complement Short Read Sequencing (SRS) technologies for routine whole genome sequencing. LRS platforms produce DNA fragment reads, from 10^3^ to 10^6^ bases, allowing the resolution of numerous uncertainties left by SRS reads for genome reconstruction and analysis. In particular, LRS characterizes long and complex structural variants undetected by SRS due to short read length. Furthermore, assemblies produced with LRS reads are considerably more contiguous than with SRS while spanning previously inaccessible telomeric and centromeric regions. However, a major challenge to LRS reads adoption is their much higher error rate than SRS of up to 15%, introducing obstacles in downstream analysis pipelines.

**Results:** We present Ratatosk, a new error correction method for erroneous long reads based on a compacted and colored de Bruijn graph built from accurate short reads. Short and long reads color paths in the graph while vertices are annotated with candidate Single Nucleotide Polymorphisms. Long reads are subsequently anchored to the graph using exact and inexact fc-mer matches to find paths corresponding to corrected sequences. We demonstrate that Ratatosk can reduce the raw error rate of Oxford Nanopore reads 6-fold on average with a median error rate as low as 0.28%. Ratatosk corrected data maintain nearly 99% accurate SNP calls and increase indel call accuracy by up to about 40% compared to the raw data. An assembly of the Ashkenazi individual HG002 created from Ratatosk corrected Oxford Nanopore reads yields a contig N50 of 43.22 Mbp and less misassemblies than an assembly created from PacBio HiFi reads.

**Availability:** https://github.com/DecodeGenetics/Ratatosk

## 1 Introduction

In Norse mythology, the squirrel Ratatöskr runs up and down the ash tree Yggdrasil, bearing envious words between the eagle at the top and the dragon at the bottom. Short read sequencing (SRS) has allowed for the accurate identification of small variants (SNPs and indels) in non-repetitive parts of the genome while long read sequencing (LRS) allows for the characterization of large and complex variations. We have designed Ratatosk to carry information between the two technologies with the hope of leveraging the benefits of both of them.

Oxford Nanopore Technologies (ONT) and Pacific Bioscience (PacBio) are LRS platforms [Logsdon et al., 2020] that produce long sequence reads ranging from 10^3^ to 10^6^ bases with an error rate up to 15% [Rang et al., 2018]. The high error rate of LRS reads is in part compensated by their lengths which increase their mapping accuracy, making LRS suitable for numerous applications in all fields of genomics. LRS used at high coverage on a few individuals [Audano et al., 2019] or low-medium coverage at population scale [Beyter et al., 2019] greatly improves the detection of Structural Variants (SVs) because the large size of ONT reads spans SVs breakpoints. Additionally, LRS reads can encompass large sections of highly repetitive regions in the human genome such as centromeres [Bzikadze and Pevzner, 2019], telomeres [Miga et al., 2020] and tandem repeats [Mitsuhashi et al., 2019]. Analyzing these regions with SRS is gruelling as the reads generally map ambiguously to multiple locations because of their limited size. Yet, centromeres play an important role in cancer genomics [Miga, 2019] while Short Tandem Repeat (STR) expansions associate with a number of genetic diseases [Kristmundsdottir et al., 2020]. LRS technologies have also enabled *de novo* haplotype-resolved assemblies with very few contig breaks [Porubsky et al., 2019]. Finally, LRS technologies overcome chemistry limitations of SRS, in particular GC bias [Chen et al., 2013] and PCR amplification artifacts [Kozarewa et al., 2009] causing uneven coverages for reads produced by Illumina platforms. Yet, the high error rate of LRS reads introduces algorithmic challenges in analyzing these data while filtering out the noise [Sedlazeck et al., 2018a]. Highly accurate LRS technologies [Wenger et al., 2019] that perform circular sequencing and generate highly accurate consensus sequences are emerging but the required resources are still prohibitive at a population scale. SRS data are therefore often used to complement to LRS data for SV breakpoint refinement [Sedlazeck et al., 2018b] and assembly polishing [Kolmogorov et al., 2019].

We present Ratatosk, a new method based on a compacted and colored de Bruijn graph for the hybrid correction of LRS reads using SRS data. Ratatosk is specifically designed to avoid over-correction with incorrect haplotypes or homologous regions as this would either remove true variants or add artificial ones. Ratatosk introduces several new features not included in other hybrid correction tools. First, SRS and LRS reads color vertices of the de Bruijn graph to highlight existing paths for the correction. Graph coloring enables pruning the search space when traversing the graph by removing chimeric paths. Second, LRS reads are anchored to the graph using both exact and inexact *k*-mer matches. The latter improves the anchoring of highly erroneous regions of the LRS reads. Third, the graph is annotated with candidate SNPs to disentangle small variations between haplotypes that are difficult to capture from erroneous LRS reads. Finally, two passes of correction are performed using SRS and LRS reads separately to take advantage of all data available, as well as increasing *k*-mer sizes to remove errors made during the first correction pass.

The performance of LRS read error correction tools is usually evaluated by the error rate, genome coverage and different assembly metrics of the corrected reads [Marchet et al., 2020, Morisse et al., 2020] but there has been little investigation of the accuracy of variant calls on the corrected data. We demonstrate that Ratatosk can reduce the raw error rate of ONT reads 6-fold on average with a median error rate as low as 0.28%. Ratatosk corrected data maintain nearly 99 % accurate SNP calls and substantially increase indel calls accuracy to nearly 92 % compared to the raw data. An assembly of the Ashkenazi individual HG002 [Zook et al., 2016] created from Ratatosk corrected ONT reads yields a contig N50 of 43.22 Mbp and less misassemblies than an assembly created from PacBio HiFi reads.

### 1.1 Previous work

Methods for LRS reads correction belong to one of two categories: self-correction or hybrid correction. Self-correction methods refine the reads using information from the set of LRS reads alone while hybrid correction methods use information from a set of SRS reads originating from the same individuals. Overall, hybrid correction methods have been shown to outperform self-correction methods in terms of error rate and compute resource usage. However, a recurrent issue with most error correction methods is that they do not retain the phasing of the reads, hence limiting the usage of corrected data to mixed-haplotype assembly. We provide here a short overview of hybrid correction methods and refer to the reviews of Zhang et al., 2019, Fu et al., 2019 and Morisse et al., 2020 for more details about self-correction methods.

Most hybrid correction methods use a de Bruijn graph built from SRS reads as an index for the correction of LRS reads. The de Bruijn graph has been extensively used as a data structure for genome assembly [Pevzner et al., 2001, Idury and Waterman, 1995]. LoRDEC [Salmela and Rivals, 2014] builds a de Bruijn graph from SRS reads and anchors LRS reads to the graph. Subsequences that do not anchor to the graph are then corrected: paths which are similar to the uncorrected subsequences are extracted and used for correction. Jabba [Miclotte et al., 2016] is similar to LoRDEC besides that SRS reads are self-corrected before graph construction and LRS reads are anchored to the graph using Maximum Exact Matches to enable different *k*-mer lengths during correction. HG-CoLoR [Morisse et al., 2018] also uses self-corrected SRS reads and aligns them to the LRS reads to find overlaps. These overlaps anchor the reads onto a variable-order de Bruijn graph allowing for multiple *k*-mer lengths. Finally, FMLRC [Wang et al., 2018] indexes the de Bruijn graph using a multi-string Burrows-Wheeler Transform of the SRS reads. This representation is lightweight in memory, enables multiple *k*-mer lengths and stores implicitly *k*-mer frequencies. FMLRC has two passes of correction, one using a short *k*-mer and one using a long *k*-mer in order to simplify the graph for high complexity regions to correct. Unlike the above tools, CoLoRMap [Haghshenas et al., 2015] constructs a weighted alignment graph from the mapping of the SRS reads to the LRS reads. The mapping provides paths in the graph that maximize the similarity with the subsequences to correct. CoLoRMap takes advantage of the paired-end information to leap over regions of LRS reads where no SRS reads map. We refer to the reviews of Zhang et al., 2019, Fu et al., 2019 and Morisse et al., 2020 for further information.

## 2 Results

We evaluated Ratatosk using our reference-guided preprocessing on a set of 4 Icelandic trios from deCODE genetics [Jonsson et al., 2017] and the Ashkenazi individual HG002 from Genome In A Bottle [Zook et al., 2016]. Each genome was sequenced with both Illumina and ONT platforms. Two HG002 data sets basecalled with Guppy 3.6 and Guppy 3.3 were employed as Guppy 3.6 significantly improves the raw read accuracy over Guppy 3.3. Genome coverage and N50 metrics are reported in Table 1 for the ONT reads. The short reads used are Illumina paired-end reads of length 151 bases with a mean coverage of 42x in the Icelandic trios and 61x in the HG002 data set. The Ratatosk corrected ONT reads were subsequently compared to the raw and FMLRC [Wang et al., 2018] corrected reads. The reviews of Zhang et al., 2019 and Fu et al., 2019 highlight FMLRC as one of the correction tools with the best overall performance among hybrid methods. Time and memory usage for Ratatosk and FMLRC are reported in Appendix, Section D. All ONT reads were subsequently aligned to the reference human genome GRCh38.p13 with minimap2 [Li, 2018] using the default ONT setting for further analysis.

**Table 1:** Genome coverage and N50 for the ONT reads of the child (C), father (F) and mother (M) in 4 Icelandic trios and the child in two HG002 data sets.

|  | Guppy version | Coverage |  |  | N50 |  |  |
| --- | --- | --- | --- | --- | --- | --- | --- |
|  |  | C | F | M | C | F | M |
| T1 | 3.3 | 63.68 | 50.94 | 64.74 | 20,353 | 24,093 | 23,528 |
| T2 | 3.3 | 55.68 | 67.95 | 70.46 | 25,767 | 22,496 | 20,439 |
| T3 | 3.3 | 69.50 | 57.05 | 56.62 | 24,047 | 27,787 | 26,856 |
| T4 | 3.3 | 57.07 | 57.40 | 64.28 | 23,111 | 15,234 | 26,634 |
| HG002-1 | 3.6 | 46.72 | NA | NA | 52,311 | NA | NA |
| HG002-2 | 3.3 | 108.10 | NA | NA | 22,260 | NA | NA |

### 2.1 Error rate

Tables 2a and 2b show the error rates for the ONT reads corrected by Ratatosk and FMLRC as well as the raw ONT reads. The mean error rate of the Ratatosk reads is about 1.9 times lower than the FMLRC reads and about 6.3 times lower than the raw reads. On the HG002 data sets, 50 % of the Ratatosk reads have an error rate of 0.28 % or below. This is up to 29 times lower than the raw reads and up to 3.5 times lower than the FMLRC reads. Details on the error rate calculations are given in Appendix, Section B.

We also report in Figure 1 the number of supplementary alignments and the ratio of ambiguous bases as metrics for read quality. Supplementary alignments occur when an alignment cannot be represented as a single linear alignment [Li et al., 2009] but instead, as a set of linear alignments where the alignment with the greatest alignment score is selected as *primary* and the others as *supplementary.* The presence of supplementary alignments might indicate an SV large enough for the aligner to abandon mapping the read with a single linear alignment. Supplementary alignments might also indicate that the read has been partially over-corrected. Finally, ambiguous bases are bases from reads which do not align in the extremities of primary alignments *(soft-clipping*) but do align in at least one distant supplementary alignment of the same reads. The ratio of ambiguous bases measure the proportion of read bases mapping ambiguously because of chimeric reads [Marijon et al., 2020] or over-correction. More details are given in Appendix, Section C.

**Figure 1:**
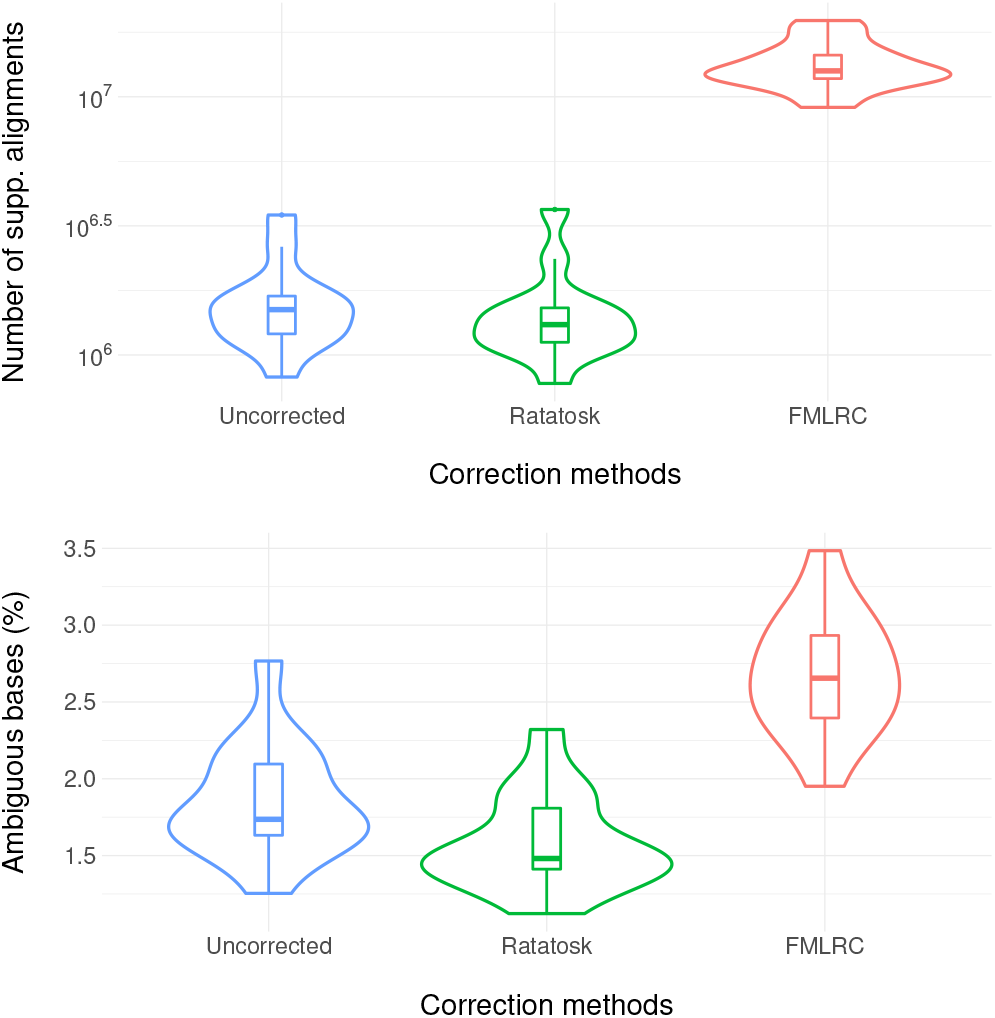
Number of supplementary alignments and ratio of ambiguous bases for the Icelandic trios and the HG002 data sets.

As shown in Figure 1, Ratatosk slightly decreases on average the number of supplementary alignments and the ratio of ambiguous bases compared to the raw reads. On the other hand, FMLRC increases the number of supplementary alignments by a factor 8.28 and increases the ratio of ambiguous bases by a factor 1.44. This suggests that Ratatosk can correct soft-clipped bases and chimeric reads while FMLRC is prone to over-correction.

### 2.2 Variant calling

There is a limited number of tools that can perform small variant calling on corrected LRS reads. Clair [Luo et al., 2020] and DeepVariant [Poplin et al., 2018] are machine learning based and can train a model given a training set of input reads. We use Clair for our evaluations as DeepVariant was several times slower to train on the raw ONT reads. Longshot [Edge and Bansal, 2019] was not used as it does not call indels while Medaka [Oxford Nanopore Technologies, accessed June 10th 2020, 2019] uses an error model specific to the raw ONT reads and hence, could not be applied to corrected data. A model was trained with Clair on the raw, Ratatosk and FMLRC ONT reads from HG002-1 using the truth set of variants less than 50 bases long in the high confidence regions (see Appendix, Section F). The different models generated for each type of input long reads were then used to call small variants on all genomes and variant calls were subsequently evaluated using rtg-tools [Krusche et al., 2019].

Given a variant truth set, rtg-tools automatically computes an optimal quality score threshold for the variant calls. Table 3a shows the variant calls accuracy for HG002-1 for which low quality variants below the optimal threshold are filtered out. On the other hand, Table 3b illustrates a standard setting for which all variants with the `FILTER` field set to `PASS` in the VCF files are used. With quality score filtering, SNP calls are nearly 99 % accurate for the raw and Ratatosk reads with a slight accuracy decrease in the SNPs called from the FMLRC reads. This demonstrates that SNPs are accurately represented in the raw reads and Ratatosk captures well the SNP candidates in the correction. However, indels are poorly represented in the raw reads and Ratatosk increases the indel calls accuracy by up to 40 % compared to the raw reads. When no filtering is applied, the difference of indel calls accuracy between raw and corrected reads is staggering. Indeed, the indel calls accuracy of raw reads shrinks to 19 % because a larger number of false positive indels are called compared to the filtered calls. Indel call accuracy from the FMLRC reads decreases to 73 % while indels called from the Ratatosk reads decline only to 91.29% accuracy, a 0.5 % reduction compared to the filtered indel calls.

**Table 2:**
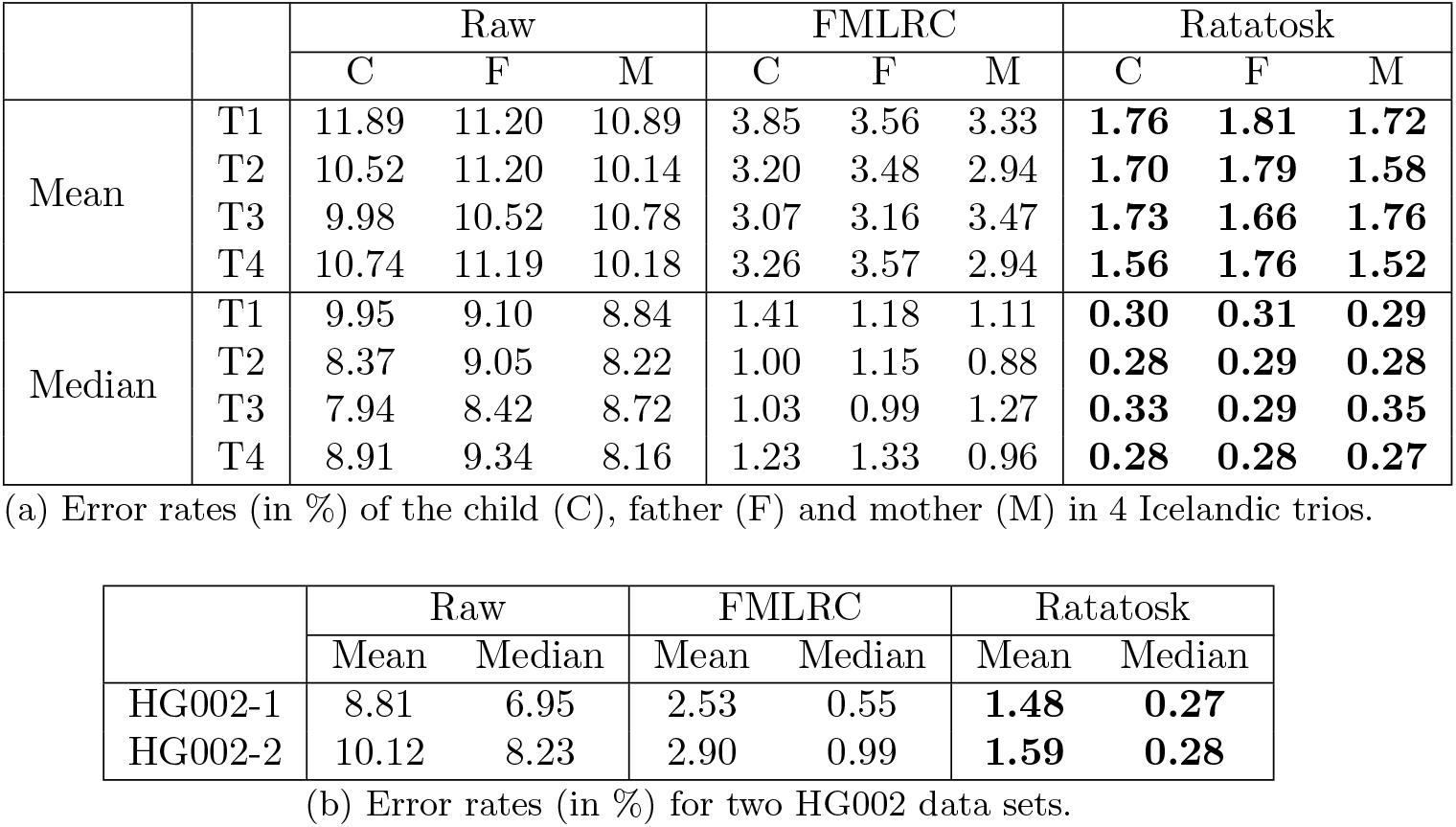
Error rates (in %) for the raw and corrected ONT reads in 4 Icelandic trios and two HG002 data sets. Best results are highlighted.

**Table 3:**
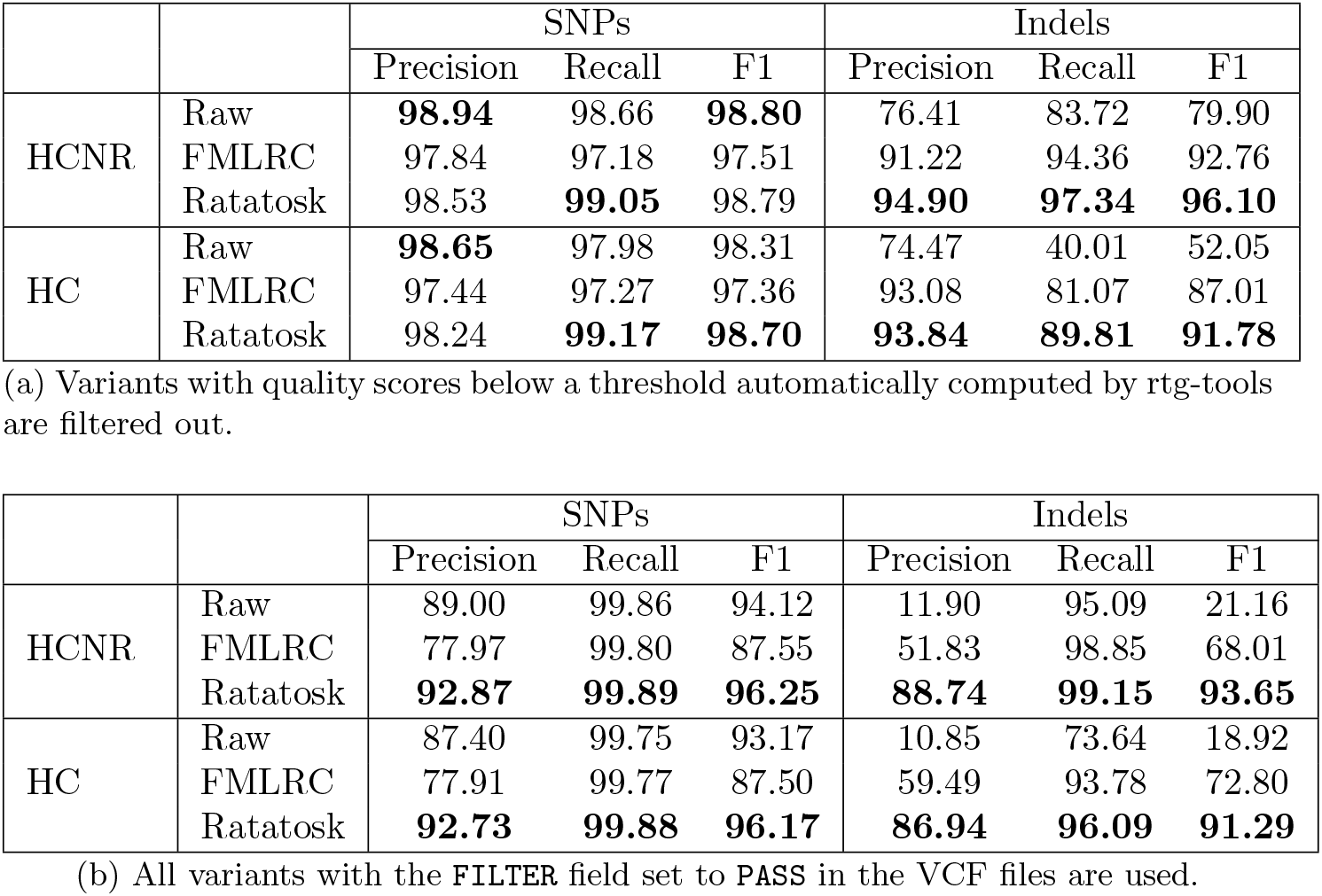
Small variant calls accuracy (in %) for the ONT reads from HG002-1 in high confidence (HC) and high confidence non repetitive (HCNR) regions. Best results are highlighted.

No variant truth set is available for the Icelandic trios so Mendelian inheritance concordance was measured by rtg-tools instead, as shown in Table 4. Note that because the trios were basecalled with Guppy 3.3, another model was trained using HG002-2 for the raw reads and FMLRC corrected reads. Overall, small variants calls from Ratatosk reads are the most consistent with the calls from each parents and both parents across all trios.

**Table 4:** Mendelian concordance (in %) of small variants called on the ONT reads of 4 children from Icelandic trios with respect to the variant calls from their father (F), mother (M) and both parents (F+M). All variants with the `FILTER` field set to `PASS` in the VCF files are used by rtg-tools. Best results are highlighted.

|  | Raw |  |  | FMLRC |  |  | Ratatosk |  |  |
| --- | --- | --- | --- | --- | --- | --- | --- | --- | --- |
|  | F | M | F+M | F | M | F+M | F | M | F+M |
| T1 | 98.87 | 98.92 | 93.04 | 98.89 | 98.96 | 95.62 | **99.49** | **99.48** | **98.03** |
| T2 | 98.90 | 98.78 | 93.05 | 99.01 | 99.00 | 95.89 | **99.50** | **99.49** | **97.70** |
| T3 | 98.86 | **98.92** | 93.70 | 98.67 | 98.57 | 95.23 | **98.94** | 98.52 | **96.25** |
| T4 | 98.80 | 98.98 | 93.39 | 98.95 | 99.12 | 96.02 | **99.51** | **99.54** | **98.21** |

### 2.3 Assembly

The Ratatosk corrected reads of HG002-1 were assembled using Flye 2.7.1 [Kolmogorov et al., 2019] and the produced contigs were evaluated with QUAST 5.0.2 [Gurevich et al., 2013] and Merqury [Rhie et al., 2020]. We compared the Flye assembly of Ratatosk corrected reads to a recent assembly made from PacBio HiFi reads with HiCanu [Nurk et al., 2020] and the reference assembly Ash1 v1.7 [Shumate et al., 2020] made from Illumina, ONT and PacBio HiFi reads assembled with MaSuRCA [Zimin et al., 2013]. The Fly/Ratatosk and HiFi/HiCanu assemblies were post-process with purge_dups [Guan et al., 2020] to exclude allelic contigs from the assemblies. Misassemblies reported by QUAST were filtered to exclude errors in known SV and segmental duplication sites as well as centromeric regions using a script from HELEN [Shafin et al., 2020]. The quality value represents a log-scaled probability of error for the consensus basecalls while the *k*-mer completeness measures the proportion of *k*-mers shared between the assembly and an accurate SRS data set from the same individual.

As shown in Table 5, the Flye assembly of the Ratatosk reads is competitive with other high quality LRS assemblies and outperforms them on several metrics. In particular, the Ratatosk/Flye assembly has the largest NG50 and NGA50, the lowest number of contigs and the smallest number of misassemblies. However, the HiFi/HiCanu assembly displays the best quality value, likely due to the high accuracy of HiFi reads. While all assemblies have a similar *k*-mer completeness, the Ash1 reference assembly has the best reference genome GRCh38 coverage. However, 1.96 % of the Ash1 assembly is derived from the reference genome GRCh38. Overall, these results demonstrate that the correction performed by Ratatosk is suited for producing highly contiguous assemblies of quality with very few errors.

**Table 5:**
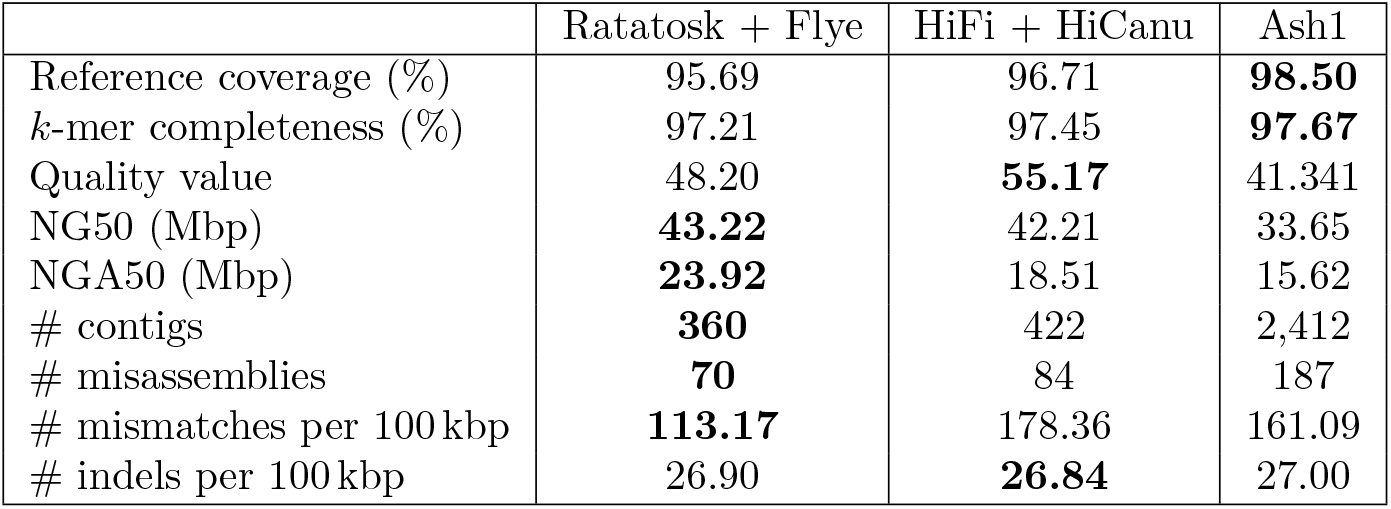
HG002-1 assembly metrics for Ratatosk corrected ONT reads assembled with Flye, PacBio HiFi reads assembled with HiCanu and the Ash1 reference assembly. Misassemblies are filtered to exclude errors in known SV and segmental duplication sites as well as centromeric regions. All metrics are computed by QUAST except *k*-mer completeness and quality value which are computed by Merqury. Best results are highlighted.

## 3 Conclusion

We present Ratatosk, a hybrid error correction tool for long and erroneous reads designed for accurate variant calling and assembly. Ratatosk uses short and long reads to color paths in a compacted de Bruijn graph index and annotate vertices with candidate Single Nucleotide Polymorphisms. We demonstrate on several data sets that Ratatosk decreases the error rate 6-fold and reduces the number of ambiguously mapped bases in the reads. SNPs calls on Ratatosk corrected reads are nearly 99 % accurate and indel calls accuracy is up to 40 % higher compared to the uncorrected reads. Furthermore, variants calls obtained from 4 corrected trios are highly concordant. Finally, we show that Ratatosk corrected data enable highly contiguous assembly with fewer errors compared to other assemblies made from accurate long reads. Future work includes trio-based correction, additional correction passes and an enhanced path coloring of the graph to enable a better correction in repetitive regions.

## 4 Methods

Section 4.1 details the concepts and data structures that will be used throughout this paper. Section 4.2 describes how the main index is built and preprocessed for correction. Sections 4.3 and 4.4 overview the methods used during the first and second correction passes, respectively.

### 4.1 Definitions

A string *s* is a sequence of symbols drawn from an alphabet 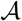. The length of *s* is denoted by |*s*|. A substring of *s* is a string in *s* with a start position *i*, a length *l* and is denoted by *s*(*i,l*). Let 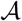 be the DNA alphabet 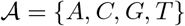 for which (*A,T*) and (*C, G*) are complementing pairs. The reverse-complemented string 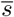 is the reverse sequence of complemented symbols in *s*. The canonical string *ŝ* is the lexicographically smallest of *s* and its reversecomplement 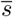. A de Bruijn graph (dBG) is a bi-directed graph *G* = (*V, E*) in which each vertex *v* ∈ *V* represents a *k*-mer and its reverse-complement. Only the canonical *k*-mer of each vertex is stored in *G*. A directed edge *ϵ* ∈ *E* from vertex *v* to vertex *v’* representing *k*-mers *x* and *x’*, respectively, exists if and only if *x*(2, *k* – 1) = *x*’(1,*k* – 1). Each edge *e* is labeled with the orientation of the *k*-mers *x* and *x*’ they connect: 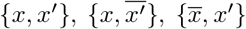 or 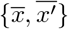. Each *k*-mer x has 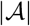 possible successors *x*(2, *k* – 1) ⊙ *a* and 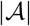 possible predecessors *a* ⊙ *x*(1, *k* – 1) in *G* with 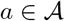 and ⊙ as the concatenation operator. The number of *k*-mers in *G* is denoted |*G*|. A path in the graph is a sequence of connected vertices *P* = (*v*_1_,…, *v_m_*). Path *P* is said *non-branching* if it is composed of vertices having an in- and out-degree of one with exception of the head vertex *v*_1_ which can have more than one incoming edge and the tail vertex *v_m_* which can have more than one outgoing edge. A non-branching path is maximal if it cannot be extended in the graph without branching. A compacted de Bruijn graph (cdBG) merges all maximal non-branching paths *P* from the dBG into single vertices, called *unitigs,* representing substrings of length |*P*| + *k* — 1. A simplified dBG and its compacted representation are illustrated in Figures 2a and 2b. A colored de Bruijn graph is a graph *G* = (*V, E, C*) in which (*V, E*) is a dBG and *C* is a set of colors such that each vertex *v ∈ V* maps to a subset of *C*. We extend the definition of a cdBG to a compacted and colored de Bruijn Graph (ccdBG) where (*V, E*) is a cDBG, so the vertices represent unitigs, and each *k*-mer of a unitig maps to a subset of *C*.

**Figure 2:**
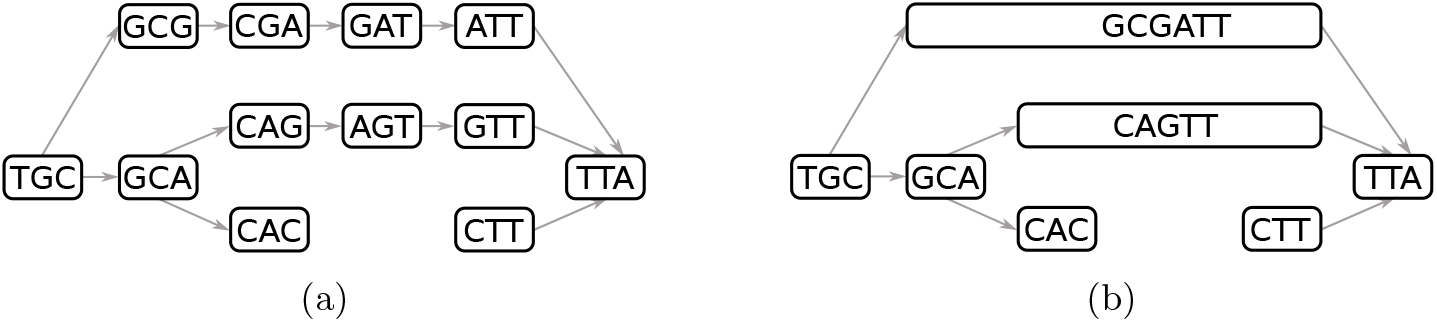
A de Bruijn graph in (a) and its compacted counterpart in (b) using 3-mers. For simplicity, the de Bruijn graph is directed and reverse-complements are not considered.

### 4.2 Graph construction and preprocessing

Ratatosk takes as input a set 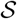 of paired SRS reads and a set 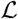 of LRS reads. A cdBG is built from 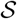 to correct the reads in 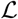 using two correction passes as shown in Figure 3.

**Figure 3:**
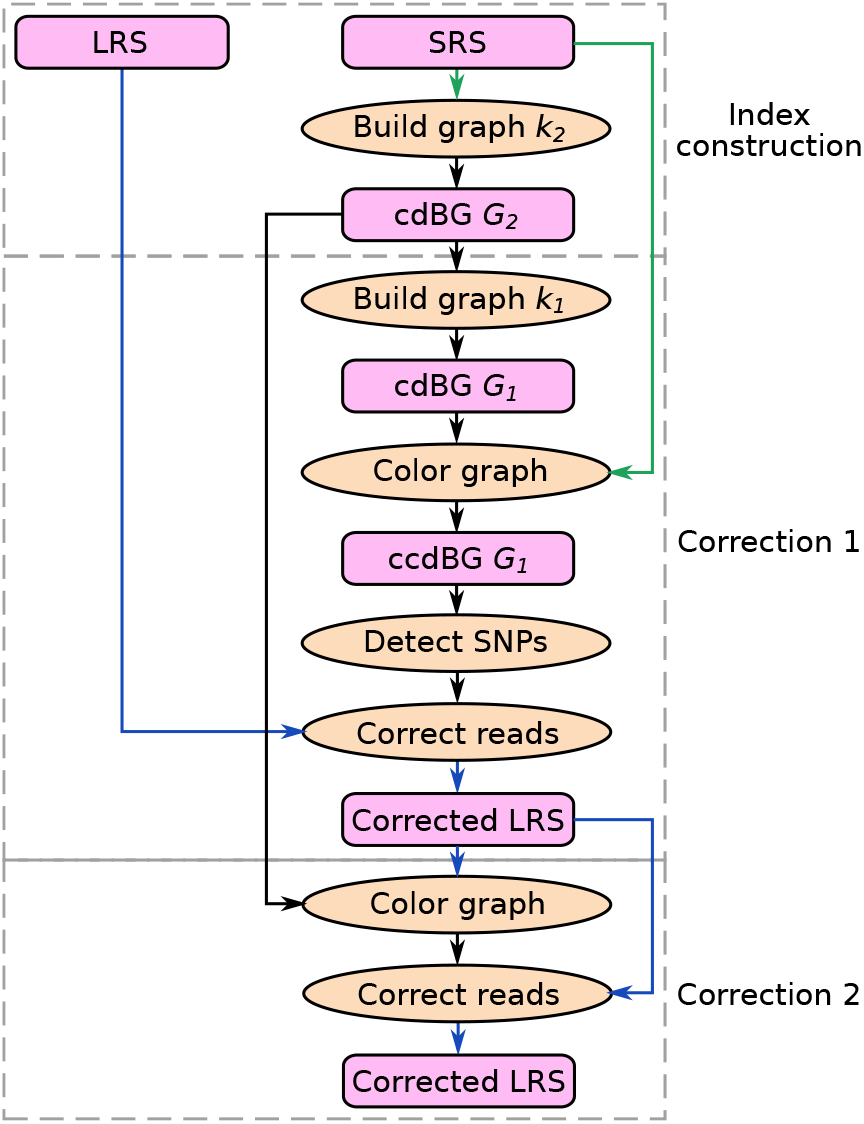
Ratatosk performs two passes of correction, each using a different *k*-mer size for the graph construction and a different type of reads for the graph coloring. LRS reads are shown in blue and SRS reads in green.

#### 4.2.1 Graph construction

Using different *k*-mer lengths in the graph built from 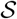 has been shown to improve the correction of 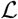 [Miclotte et al., 2016, Morisse et al., 2018, Wang et al., 2018]: A short *k*-mer is ideal for finding matches between LRS reads and the graph while unitigs built with long *k*-mers have a better contiguity. In order to combine the advantages of short and long *k*-mers, Ratatosk uses two *k*-mer lengths *k*_1_ and *k*_2_ with *k*_2_ ≥ 2*k*_1_.

First, a cdBG *G*_2_ is built with the long *k*_2_-mers of 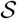 using the Bifrost graph engine [Holley and Melsted, 2019]. By default, all *k*_2_-mers occurring exactly once in 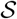 are assumed to contain a sequencing error and are discarded from the graph construction. Subsequently, a cdBG *G*_1_ is built from the short *k*_1_-mers of the unitigs in *G*_2_. Graph *G*_1_ is used for the first correction pass while graph *G*_2_ is later used in the second correction pass (Figure 3, Section 4.4).

#### 4.2.2 Graph coloring

Graph *G*_1_ is turned into a ccdBG by coloring its unitigs with the read pairs from 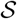 with which they share at least one *k*_1_-mer, as shown in Figure 4. Given 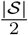 SRS read pairs in input, each pair is identified by a color ranging from 1 to 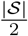. Ratatosk uses hashing to ensure that two reads from the same pair get the same color. Each read is initially given an identifier which is the hash of the read name and each identifier is associated with a color. To simplify the coloring and curb its running time, pair identifiers are not guaranteed to be unique such that multiple read pairs might hold the same identifier, hence the same color, because their read names hash to the same value. These collisions are random and are expected to have little impact on the correction.

**Figure 4:**
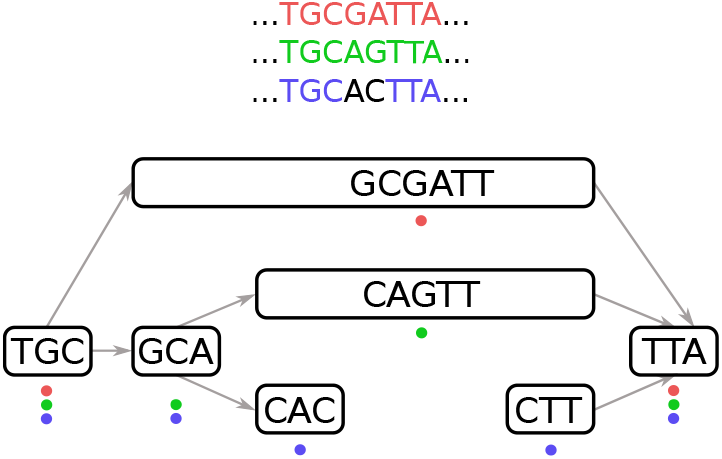
Graph coloring with three colors. Each color represents a read sharing at least one *k*-mer with a unitig.

Coloring unitigs with read pairs is similar to *partitions* in the guided de Bruijn graph [Holley et al., 2017] and *links* in the Linked de Bruijn graph [Turner et al., 2018]. Ratatosk enables a memory efficient graph coloring by using two techniques. First, it discards *similar* read pairs from the coloring as they represent redundant information to index, especially in the case of high coverage SRS data. Second, Ratatosk compacts the set of colors assigned to each unitig by using consecutive color values rather than random values. Consecutive color values can be compacted in memory using Run Length Encoding, delta encoding and compacted bitmaps [Chambi et al., 2016].

The graph coloring objective is for each read pair 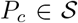 to color a set of unitigs 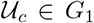 with color *c*. Each unitig 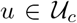 shares at least one *k*_1_-mer with *P_c_* as shown in Figure 4. We devised a probabilistic algorithm which colors the graph while discarding similar read pairs and assigning consecutive color values to unitigs in a greedy fashion. The algorithm performs two steps: A partial coloring of the graph is initially made for read pair filtering and color compaction purposes before completing a full graph coloring. In the partial coloring, each read pair *P_c_* will only color the longest unitig 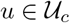. This unitig acts as a centroid in the graph and it is expected that read pairs similar to *P_c_* will color the same unitig. Hence, color *c* and a hash 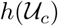 are assigned to unitig *u* unless there exist another pair similar to *P_c_* which has already been assigned to *u*. Two read pairs *P_c_* and 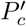 are similar if they have the same unitig set hash, i.e, *c* ≠ *c*’ but 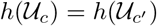. In which case, *P_c_* is discarded and will not be used for the full graph coloring. Next, read pairs are given new colors by assigning, when possible, consecutive color values to pairs clustering to the same unitigs. A final graph coloring is then performed using read pairs which are not discarded and their new color values. The mean *k*_1_-mer coverage of unitigs is also computed during the final graph coloring.

After coloring the graph, all unitigs with a mean *k*_1_-mer coverage lower than a pre-defined threshold *T_min_* (see Appendix, Section A) are removed from the graph. In order to limit even further the memory usage of Ratatosk, unitigs having a mean *k*_1_-mer coverage greater than *T_max_* have their colors discarded. These unitigs are usually *k*_1_-mers with a low sequence entropy such as poly-{*A, C, G, T*} or *k*_1_-mers occurring within STRs. They are typically located within highly branching subgraphs and their colors provide little guidance information while severely impacting the running time.

#### 4.2.3 Candidate SNP annotation

While most *de novo* detection methods for SNPs, indels and SVs are based on the analysis of graph *bubbles* [Onodera et al., 2013, Peterlongo et al., 2017, Paten et al., 2018, Garrison et al., 2018], Ratatosk uses instead a simple but fast string matching method to annotate vertices in the graph containing one or more candidate SNPs. For each *k*_1_-mer *x* in unitigs, the graph is queried for all *k*_1_-mers having a Hamming distance of 1 with *x*. Let *x* = *u*(*p, k*_1_) and *x*’ = *u*’(*p*’,*k*_1_) be *k*_1_-mers from unitigs *u* and *u*’, respectively, that differ by exactly one substitution at position *i* < *k*_1_. Unitigs *u* and *u*’ are then annotated at position *p* + *i* and *p*’ + *i*, respectively, with a IUPAC symbol representing the substitution. For example, symbol R would be assigned to position 3 in unitigs GCGATT and GCA of Figure 4 to represent an A/G substitution.

### 4.3 First correction pass

The following section describes how LRS reads are anchored to the ccdBG and the methods used to correct non-anchored regions of the LRS reads.

#### 4.3.1 Read Anchoring

We define *solid* and *weak k*-mers similarly as defined in LoRDEC and introduce the definition of *near solid k*-mers:

- solid *k*-mer: exact length *k* substring match between a long read and a unitig from the graph.
- near solid *k*-mer: inexact length *k* substring match between a long read and a unitig from the graph with one base substitution or indel.
- weak *k*-mer: length *k* substring of a long read which is neither a solid *k*-mer nor a near solid *k*-mer.

We define two types of regions in a long read:

- solid region: a region of a long read composed only of solid *k*-mers.
- non-solid region: a region of a long read composed of weak or near solid *k*-mers.

A solid or near solid *k*-mer is also called a *match*. A match between long read *r* at position *p_r_* and unitig *u* at position *p_u_* is denoted *m* = 〈*p_r_,r,p_u_,u*〉. A match *m* is *unique* if it is the only match at position *p_r_* in *r*. A *k*-mer has at most one solid match in *G*_1_ but can have multiple near solid matches in *G*_1_. Note that solid and non-solid regions can overlap by *k* — 1 bases. All non-solid regions are surrounded by two solid regions with the exception of non-solid regions at the start and end of LRS reads.

#### 4.3.2 Delimiting non-solid regions

Each read 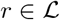 is corrected independently, allowing multiple threads to correct LRS reads in parallel. The graph is queried for each *k*_1_-mer of *r*, resulting in a list of solid matches *M_s_* and a list of near solid matches *M_n_*, both sorted by ascending match position *p_r_* in *r*. Only unique near solid matches (UNSM) are kept in *M_n_* to prevent anchoring *r* on a SNP or indel from an incorrect allele. Furthermore, a *k*_1_-mer which is both a solid match and a near solid match is considered solid and its near solid matches are discarded from *M_n_*.

Non-solid regions of *r* are detected by finding all pairs of successive solid matches *m_s_,m_t_* ∈ *M_s_* for which 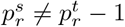 with the exception of non-solid regions at the extremities of *r*. The first match *m_s_* of the pair is referred to as the *source* match and the second match *m_t_* of the pair is referred to as the *target* match. The length of the non-solid region to correct is then 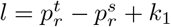. It includes 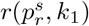 which is the last solid *k*_1_-mer from the source solid region and 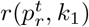 which is the first solid *k*_1_-mer from the target solid region as illustrated in Figure 5. If a read starts with a non-solid region, that region has no source match and hence starts on the first position of the read. Similarly, if a read ends with a non-solid region, that region has no target match and hence ends on the last position of the read.

**Figure 5:**
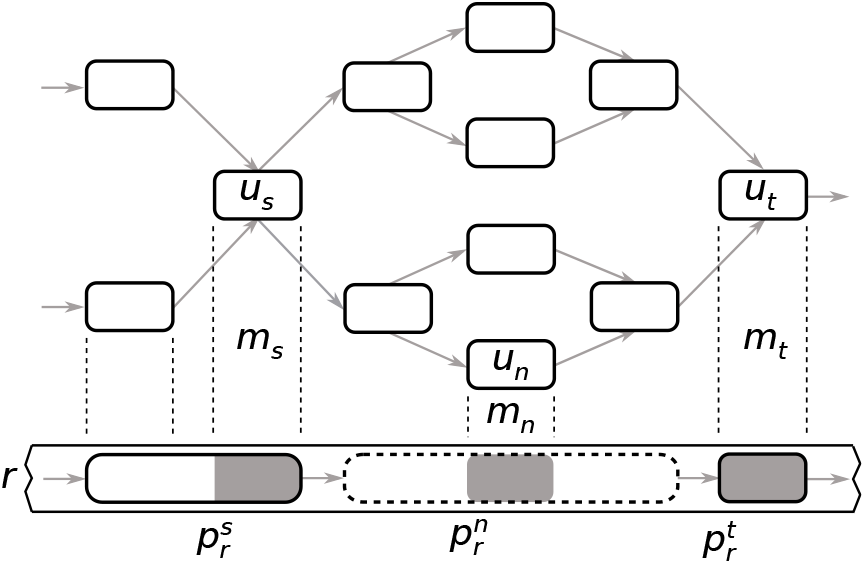
Example of a long read *r* anchored on a ccdBG. A section of *r* is shown at the bottom with two solid regions (non-dashed boxes at the extremities) surrounding a non-solid region (dashed line box at the center). The grey areas of the solid regions show the source and target matches between the long read and the graph. The grey area of the non-solid region shows a near solid match. For simplicity, colors are not shown.

#### 4.3.3 Traversing the graph

In order to correct a non-solid region, Ratatosk attempts to extract one path in the graph connecting unitig *u_s_* of the source match to unitig *u_t_* of the target match. Since the length *l* of the non-solid region to correct is known, we assume that the corrected path between *u_s_* and *u_t_* has minimum sequence length 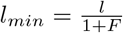 bases and maximum sequence length *l_max_* = *l* · (1 + *F*) bases where *F* is an upper-bound of the error rate in the long read (see Appendix, Section A). Ratatosk uses two greedy techniques to guide the traversal in the graph and prune the search space, as shown in Figure 6.

**Figure 6:**
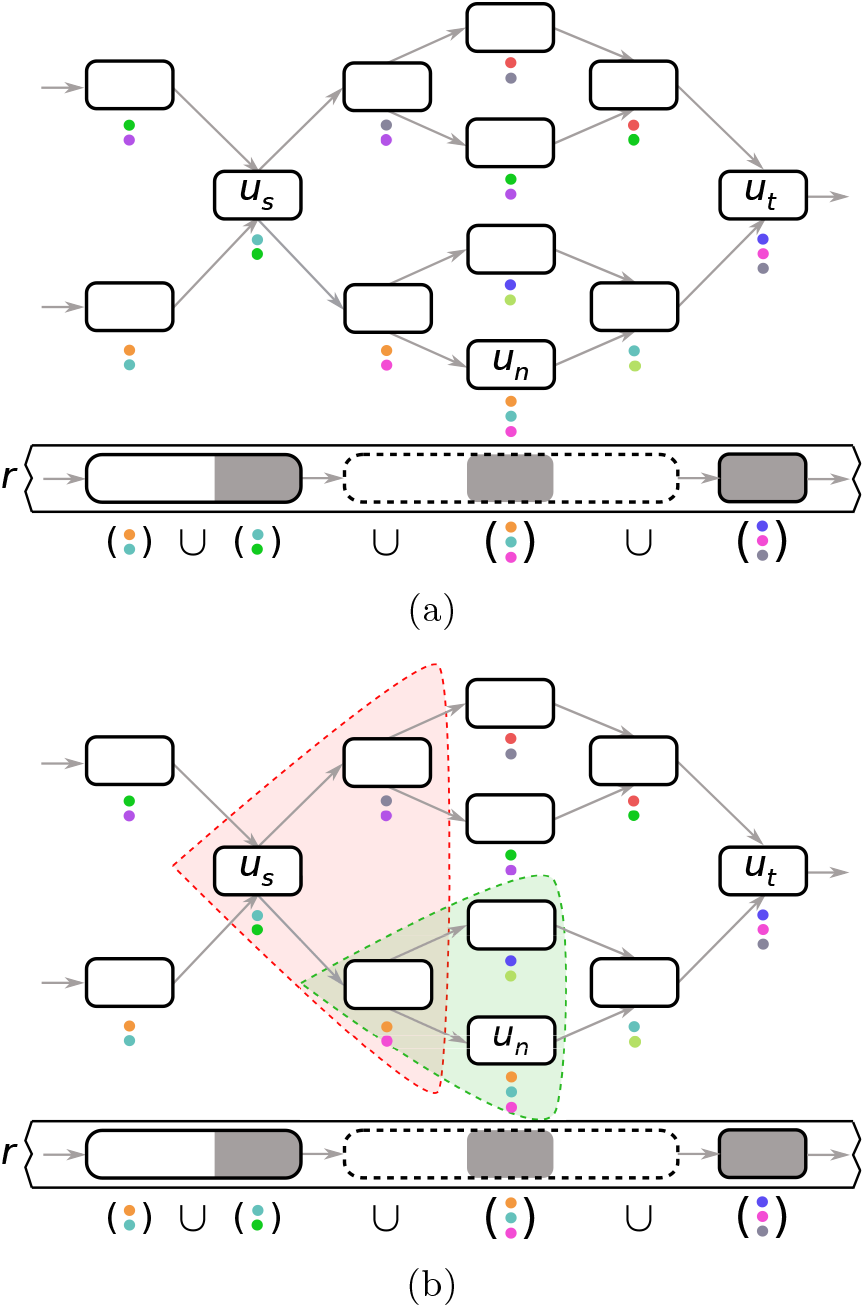
In (a), the union of colors is computed within the solid regions around the non-solid region to correct and the USNMs. This union will partially guide the graph traversal, along with the sequence similarity of the paths to the nonsolid region. In (b), a first subgraph (highlighted in red) of all paths starting at unitig *u_s_* with *P_max_* = 2 unitigs is explored for correction. The lower path is extended using the same method (shown in green) and a path connecting to *u_n_* is found.

First, rather than exploring all paths between unitigs *u_s_* and *u_t_*, Ratatosk only explores paths traversing UNSMs in the non-solid region to correct. These matches provide an anchoring in the non-solid regions as they are near exact *k*_1_-mer matches between the graph and the read to correct. Hence, paths between *u_s_* and *u_t_* which do not traverse the UNSMs are pruned because they are not good candidates for the correction. Let *m_n_* be the near solid match from *M_n_* with the smallest position 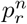 such that 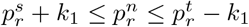. Ratatosk first attempts to extract one path connecting unitig us to unitig *u_n_* ∈ *m_n_* with a BFS traversal that only explores paths with maximum sequence length 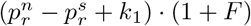 bases. The extracted path is then extended from *u_n_* to the next UNSM in *M_n_*. The process of extending the last unitig of a path to the next UNSM in *M_n_* is repeated until there are no more UNSMs to consider in *M_n_* or no path extension is possible. Finally, the graph traversal attempts to extend the path to the target unitig *u_t_*. Note that in the absence of UNSM in the non-solid region to correct, all paths connecting *u_s_* and *u_t_* with minimum sequence length *l_min_* and maximum sequence length *l_max_* are traversed.

Second, even using UNSMs to prune the search space during traversal, the subgraph between two unitigs *u_n_* and *u_n_’* from UNSMs can be very large. This is particularly true for LRS reads with a high error rate, resulting in long non-solid regions with few or no UNSMs. In order to prune the search space between *u_n_* and *u_n_’*, a greedy graph traversal is used to extract one path connecting the two unitigs. Unitig *u_n_* is first extended by visiting all paths of length *P_max_* vertices with a BFS traversal. Each traversed path is given a probability *s_P_* of being the correct path to extend and only the path with the greatest probability is extended. The path chosen for extension maximizes its sequence similarity with the non-solid region to correct. Furthermore, as colors highlight paths in the graph representing SRS reads, the path chosen for extension also maximizes its color similarity with the surrounding solid regions. Hence, before correcting a non-solid region, Ratatosk first computes the union *C* of all colors sets *C_u_* from the solid matches and UNSMs within an interval corresponding to the non-solid region start and end positions extended of *B* bases on each side, i.e,

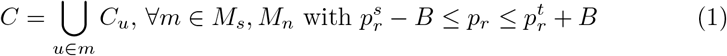

During the BFS traversal, a path probability *s_P_* is computed for each traversed path based on the number of colors the path shares with *C* and the sequence similarity of the path to the region to correct. Specifically, given a path P composed of *P_max_* unitigs and its color set 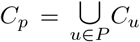, the color matching probability of *P* is 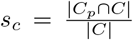 and the sequence matching probability *s_q_* is derived from the normalized edit distance of *P* to the non-solid region to correct using an infix alignment computed by the edlib tool [Šošić and Šikić, 2017]. Both probabilities are then conflated:

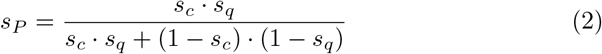

The path with the greatest probability *s_P_* is extended by starting a new graph traversal from its last unitig. The extension continues until unitig *u_n_*’ is reached or no path can be extracted as a result of a tip in the graph or extending over 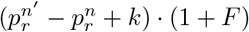 bases.

To enable a faster traversal, a local minimum number of colors *T_C_* is computed from the surrounding solid regions and the unitigs of UNSMs. Each traversed unitig *u* of a path *P* must be colored by at least *T_C_* colors of *C* such that:

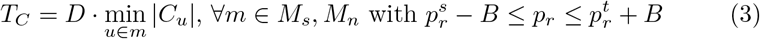

and *D* being a fixed lower bound factor (see Appendix, Section A). If the color set of a traversed unitig has less than *T_C_* colors, its path is not explored any further nor it is considered for extension.

A path extension connecting unitig *u_u_* to unitig *u_v_* might end prematurely for multiple reasons: all possible extensions end on a tip of the graph because of incomplete SRS data or insufficient color coverage in the traversed subgraph. In such a case, the extended path is completed with a gap corresponding to the non-solid subsequence to correct and the path extension resumes from unitig *u_v_*. An example of path extension with a gap is illustrated in Figure 7.

**Figure 7:**
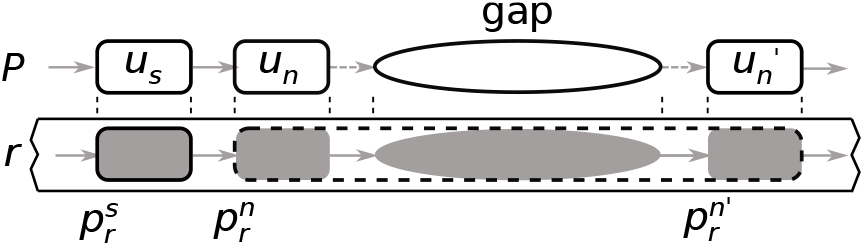
Example of a gap in a path. Path *P* is first extended until unitig *u_n_*, then a gap corresponding to a subsequence from the uncorrected read is inserted in *P* and the extension of *P* resumes from unitig *u_n_*’.

Finally, non-solid regions located on long read extremities have only one surrounding solid region. The non-solid region at the start of a long read is corrected using a backward graph traversal from *u_t_* and the one at the end of a long read is corrected with a forward graph traversal from *u_s_*. Because each of these graph traversals has no target match, any path with length *l* base such that *l_min_* ≤ l ≤ *l_max_* is returned as a candidate for correction.

#### 4.3.4 Forward and backward corrections

A candidate path for correction is *incomplete* if it contains a gap or if it does not connect to the unitig of a target match. If no path or only an incomplete path has been extracted, Ratatosk corrects the non-solid region backward, i.e., from the target match to the source match. Indeed, the forward graph traversal might have stopped prematurely for multiple reasons, one of which being that the color guidance led incorrectly to a tip in the graph. However, traversing the graph backward might lead to a different path. If both forward and backward paths are incomplete, Ratatosk merges both paths by aligning their sequences to the non-solid region using the Needleman-Wunsch algorithm (global alignment). The merged sequence is created by traversing the alignment of both forward and backward corrections at the same time and selecting subsequences in each corrections. In the case of candidate paths starting or ending a long read, all candidate paths are aligned to the non-solid region using a local alignment that does not penalize gaps at the end. The candidate path with the smallest edit distance is chosen for the correction.

#### 4.3.5 Candidate SNP correction

Heuristics used to traverse the graph as presented in Section 4.3.3 might incorrectly extend a path and lead to the erroneous correction of a non-solid region using SNPs from incorrect alleles. Once a path has been selected to correct a non-solid region, all the positions in this path matching candidate SNPs and their IUPAC symbols are known from the unitigs. Let s be the non-solid region and *s*’ its corrected counterpart. Sequence *s*’ is aligned to s and a CIGAR string is generated from the alignment. Ratatosk iterates over matching positions of the CIGAR string (symbol M) denoted *m* = 〈*s,p, s*’,*p*’〉. Note that *m* indicates that base *s*(*p*, 1) is either a match or a mismatch with base *s*’(*p*’, 1) but is not part of an insertion or deletion in the alignment. Let *M_snp_* be the set of all matches *m* = 〈*s,p, s*’,*p*’〉 for which *s*’(*p*’, 1) has an assigned IUPAC symbol in the graph indicating a candidate SNP. For each match *m* = 〈*s,p, s*’,p’〉 ∈ *M_snp_*, base *b* = *s*(*p*, 1) is compared to the IUPAC symbol associated to *b*’ = *s*’(*p*’, 1). If *b* is one of the possible bases represented by the IUPAC symbol, then *b*’ is corrected with *b*. This method enables a conservative correction of SNPs in the corrected non-solid regions by using only bases from the uncorrected nonsolid regions which are compatible with the candidate SNPs from the graph. However, this method only corrects SNPs in the matching or mismatching regions of the alignment and discards candidate SNPs located within insertions of *s*’. To overcome this issue, a match *m* ∈ *M_snp_* is said *strongly compatible* if *s*’(*p*’, 1) = *s*(*p*, 1) prior to SNP correction. A strongly compatible SNP indicates that Ratatosk is confident in the subpath that was selected to correct the region around that candidate SNP. As the strongly compatible SNP at position *p*’ is from unitig *u*’ ∈ *m*, all bases which are candidate SNPs in *u*’ are used to correct SNPs in the inserted positions of the alignment (symbol I in the CIGAR string) around position *p*’.

### 4.4 Second correction pass

In the first correction pass, Ratatosk corrected each LRS read independently from the other reads in 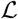. In a second correction pass, Ratatosk takes advantage of the set of corrected LRS reads as a whole. Indeed, reads corrected during the first pass might be sufficiently error-free to correct the remaining non-solid regions. Furthermore, LRS reads are at least an order of magnitude longer than SRS reads and do not need to be paired, hence offering more information to which paths to traverse in the graph. In the following, we describe the second correction pass with a highlight on the differences with the first correction pass.

Let 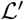 be the set of corrected LRS reads obtained from the first correction pass. First, graph *G*_2_ built from the *k*_2_-mers of 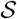 (Section 4.2.1) is loaded in memory. Compared to *G*_1_, unitigs of *G*_2_ have a better contiguity and some of the highly branching subgraphs of *G*_1_ corresponding to repetitive regions are untangled in *G*_2_. Graph coloring and candidate SNP annotation using 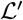 are performed as described in Sections 4.2.2 and 4.2.3, respectively. Because the reads in 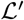 are long and still erroneous in the uncorrected regions, they are not expected to be similar and Ratatosk does not perform similar reads removal.

Reads of 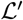 are then anchored on the graph and non-solid regions are corrected as described in Section 4.3. Parameter *B* in Equation 1 corresponds to the size of a buffer around a non-solid region where the union of unitig colors from solid and UNSMs is computed. In the first correction pass, solid regions are expected to be short and sparse because of the high error rate of LRS reads. Hence, *B* was large enough to span two SRS reads from the same pair and the gap that intersperse them in order to capture as many colors as possible. Corrected LRS reads have no gap and are much longer than SRS reads so it is expected that solid regions are much more abundant and contiguous than during the first correction pass. Distance *B* is therefore much smaller for the second pass (Appendix, Section A) which saves computation time. Furthermore, solid regions are required to be at least *B* > *k*_2_ bases long in the second pass to increase the contiguity of solid regions and provide a better anchoring on the graph.

During path selection described in Section 4.3.3, BFS traversals explored all paths of *P_max_* unitigs and a path probability was assigned to each one of them before selecting one path for extension. Traversing a fixed number of unitigs avoids a combinatorial growth of the number of explored paths, especially in complex subgraphs with short cycles that are characteristic of STRs. However, as unitigs can have any length ≥ *k*_1_, it has the disadvantage that the path probability might be computed for paths of *P_max_* unitigs with different sequence lengths. Instead, the graph traversal in the second correction pass explores paths with a minimum sequence length of *B* bases rather than a minimum number of unitigs.

Once a path *P* has at least *B* bases in its sequence, its color matching probability *s_c_* and sequence matching probability *s_q_* are computed and conflated into a path probability *s_P_*. The construction of color set *C* used in the color matching probability *s_c_* is shown in Equation 4 and only uses the intersection of colors from each side of the non-solid region, i.e, *C^s^* and *C^t^*, rather than the union (Equation 1) in order to remove erroneous colors which do not belong to this region:

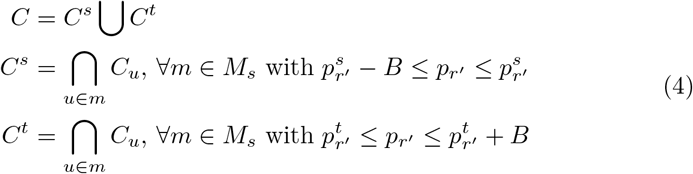

### 4.5 Reference-guided correction

While Ratatosk is a reference-free method, we propose an optional reference-guided preprocessing of the reads which is beneficial in several ways. This pipeline first maps the input SRS and LRS reads to a reference genome and then clusters the reads into *bins* corresponding to 5Mbp long regions of the reference. Each bin of SRS and LRS reads is subsequently corrected independently. The benefit is three-fold:

- Graphs *G*_1_ and *G*_2_ built from an SRS bin are much smaller and contiguous than for the entire SRS data set, hence reducing the probability of selecting an incorrect path during graph traversal.
- Computation time is reduced as the search space in each bin is much smaller than for the entire SRS data set.
- Each bin is corrected independently so the workload can be distributed in parallel over many nodes of an HPC.

However, a reference-guided preprocessing also introduces some challenges. First, it is common that reference genomes contain gaps. For example, the human genome reference GRCh38.p13 has about 161 Mbp of N bases. Second, SRS reads overlapping large insertion events are expected to be unmapped. Finally, SRS reads with poor mapping qualities map ambiguously to the reference and might be incorrectly binned.

To overcome these issues, Ratatosk detects reads from the full SRS data set 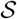 which are likely missing in each bin. Let 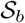 and 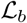 be the subset of SRS and LRS reads of a bin *b*, respectively. To begin with, cdBGs 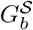 and 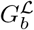 are built from the *k*_1_-mers occurring twice or more in 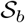 and 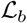, respectively. Once 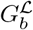 is built, its unitigs are annotated with their mean *k*_1_-mer coverage. At first, 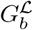 contains many more *k*_1_-mers than 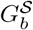 because many erroneous *k*_1_-mers from 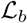 occur twice or more in the bin. To prune these erroneous *k*_1_-mers from 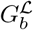, unitigs having low coverages are removed iteratively until 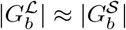. Subsequently, all reads 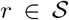 are queried: If *r* contains many *k*_1_-mers occurring in 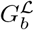 but not in 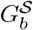, *r* is suspected to be missing from the bin and is added to 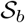.

We outline the binning and correction pipeline proposed, as illustrated in Figure 8, in the following. First, all reads from 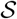 and 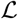 are binned into regions of 5 Mbp according to their mapping to the reference genome. Low mapping quality (< 30) and unmapped LRS reads are set aside in a bin for ambiguous long reads. Once all reads have been binned, a local correction is performed in parallel for all non-ambiguous bins and the output corrected LRS reads are merged. Note that each bin correction has access to 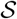 (top red arrows in Figure 8) to retrieve the missing SRS reads from the bin. Finally, the bin of ambiguous LRS reads is corrected globally using 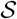. This correction is assisted by the previously corrected non-ambiguous LRS reads to enhance graph coloring during the second round of correction (bottom red arrow in Figure 8).

**Figure 8:**
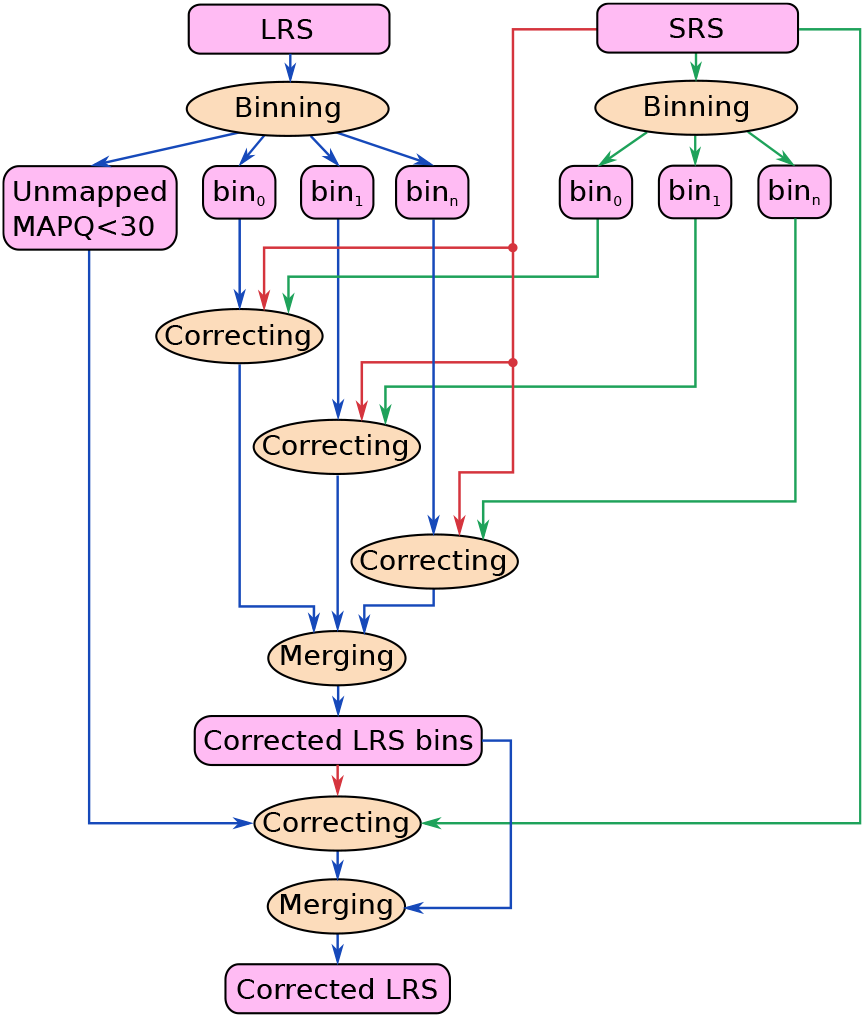
Reference-guided preprocessing of the input SRS reads (green) and LRS reads (blue). Reads are first binned and each bin is corrected independently. Unmapped or low mapping quality LRS reads are corrected using all SRS reads and all corrected LRS bins. Red arrows indicate input read sets which assist with the correction but are not corrected themselves.

## Acknowledgements

The authors would like to thank our colleagues from deCODE genetics and Amgen Inc. as well as Peter Loof Møller from Aarhus University for their helpful feedback during the development of Ratatosk. We would also like to thank Rosemary Dokos and Philipp Rescheneder from Oxford Nanopore Technologies for their feedback on Ratatosk and providing the HG002 data set. Finally, we thank all research participants who provided a biological sample to deCODE genetics and to the Genome in a Bottle Consortium.

## Author’s contributions

GH implemented the Ratatosk software. GH and BVH designed the Ratatosk algorithm with input from DB, HI, SK and HPE. GH and BVH designed the experiments. GH analyzed the data sets. GH wrote the initial version of the manuscript. All authors contributed to subsequent versions. All authors reviewed and approved the final version of the manuscript.

## A Default parameters

Graph construction and coloring:

- *k*_1_ =31
- *k*_2_ =63
- *F* = 0.25
- *T_max_*(*u*) = 512 · (|*u*| – *k* + 1) for *u* ∈ *V*

First correction pass:

- *T_min_* – 2
- *B* = 500
- *D* = 0:1
- *P_max_* = 4

Second correction pass:

- *T_min_* – 1
- *B* = 95

## B Error rate

Let *r* be an LRS read which has been aligned to a reference genome. We define the following:

- |*r*|: Number of bases in *r*
- *D_r_*: Number of deleted bases in alignment of *r* to reference
- *I_r_*: Number of inserted bases in alignment of *r* to reference
- *M_r_*: Number of mismatching bases in alignment of *r* to reference
- *S_r_*: Number of soft-clipped bases in alignment of *r* to reference

The deletion, insertion and substitution error rates of a set of LRS reads 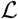 are:

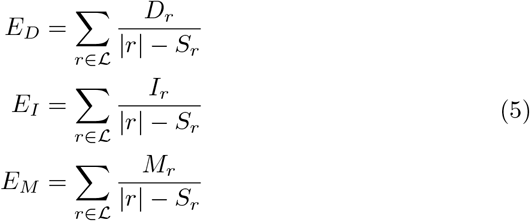

And the combined error rate of *R* is:

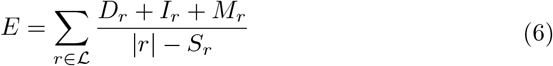

The error rates are computed from primary alignments only.

## C Ambiguous bases

Ambiguous bases are bases which soft-clip in the primary alignment but map in non-overlapping supplementary alignments to distant reference positions, such as different chromosomes, compared to the primary alignment reference position. The ratio of ambiguous bases aim to measure the proportion of over-corrected bases in reads. To avoid counting ambiguous bases in supplementary alignments of a read *r* that might correspond to a large SV event, a supplementary alignment is only used if it does not overlap the primary alignment nor another supplementary alignment of *r* with a large buffer of 1 Mbp on each side of the alignment. Algorithm 1 details the ambiguous bases ratio computation for a read represented by a primary alignment and a set of supplementary alignments.

**Algorithm 1.**
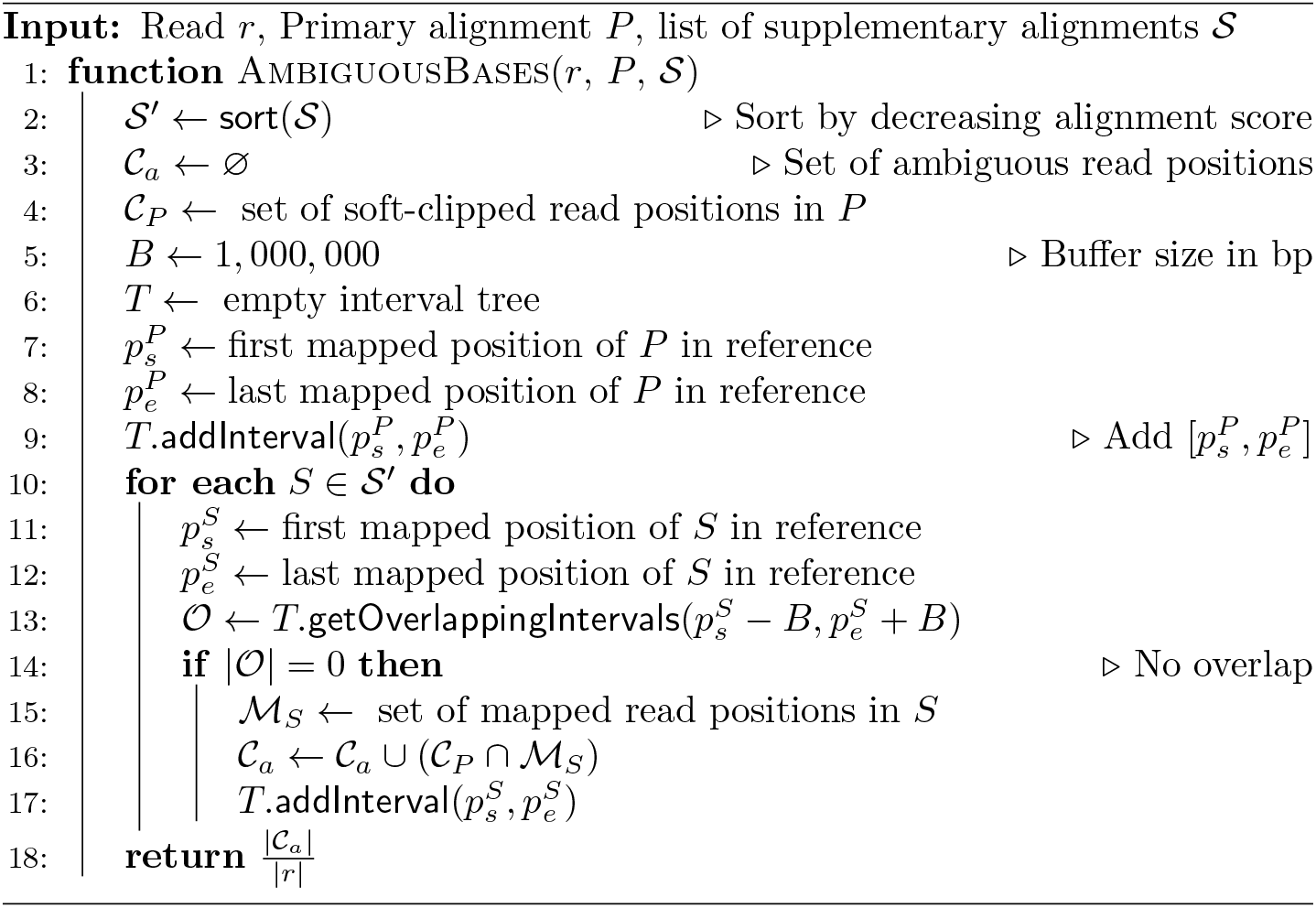
Compute ratio of ambiguous bases

## D Time and memory

|  | Time (CPU h.) |  | Peak RAM (GB) |  |
| --- | --- | --- | --- | --- |
|  | Min | Max | Min | Max |
| Ratatosk | 11,698 | 22,252 | 213.22 | 305.29 |
| FMLRC | 4,228 | 10,828 | 122.69 | 253.49 |

The reported running times do not include data preprocessing for both tools which account for a negligible amount of time and memory compared to the total running time. On average, Ratatosk is 3.13 times slower than FMLRC. However, preliminary results on the HG002-1 data set suggests that Ratatosk v0.2 is 2.57 times faster than Ratatosk v0.1 which was used in our experiments.

Ratatosk was run in parallel on several machines due to the reference-guided preprocessing of the input data while FMLRC was run on a single machine. The reported peak of memory for Ratatosk matches the correction of the ambiguous LRS bin at the end of the preprocessing pipeline as it requires to use all the input SRS data. Indeed, Ratatosk memory usage is dominated by the graph index and the SRS data coloring of its vertices during the first correction pass.

## E Tools

| Tool | Version / Commit | URL |
| --- | --- | --- |
| Ratatosk | 0.1 | https://github.com/DecodeGenetics/Ratatosk |
| FMLRC | 77dde49 | https://github.com/holtjma/fmlrc |
| minimap2 | 2.14-r883 | https://github.com/lh3/minimap2 |
| Clair | 2.0.6 | https://github.com/HKU-BAL/Clair |
| rtg | 3.10.1 | https://github.com/RealTimeGenomics/rtg-tools |
| Flye | 2.7.1 | https://github.com/fenderglass/Flye |
| QUAST | 5.0.2 | https://github.com/ablab/quast |
| Merqury | ed5918c | https://github.com/marbl/merqury |
| purge_dups | fe8dce2 | https://github.com/dfguan/purge_dups |

Script `quast_sv_extractor.py` was obtained from HELEN (https://github.com/kishwarshafin/helen/tree/master/helen/modules/python/helper).

## F HG002 Data

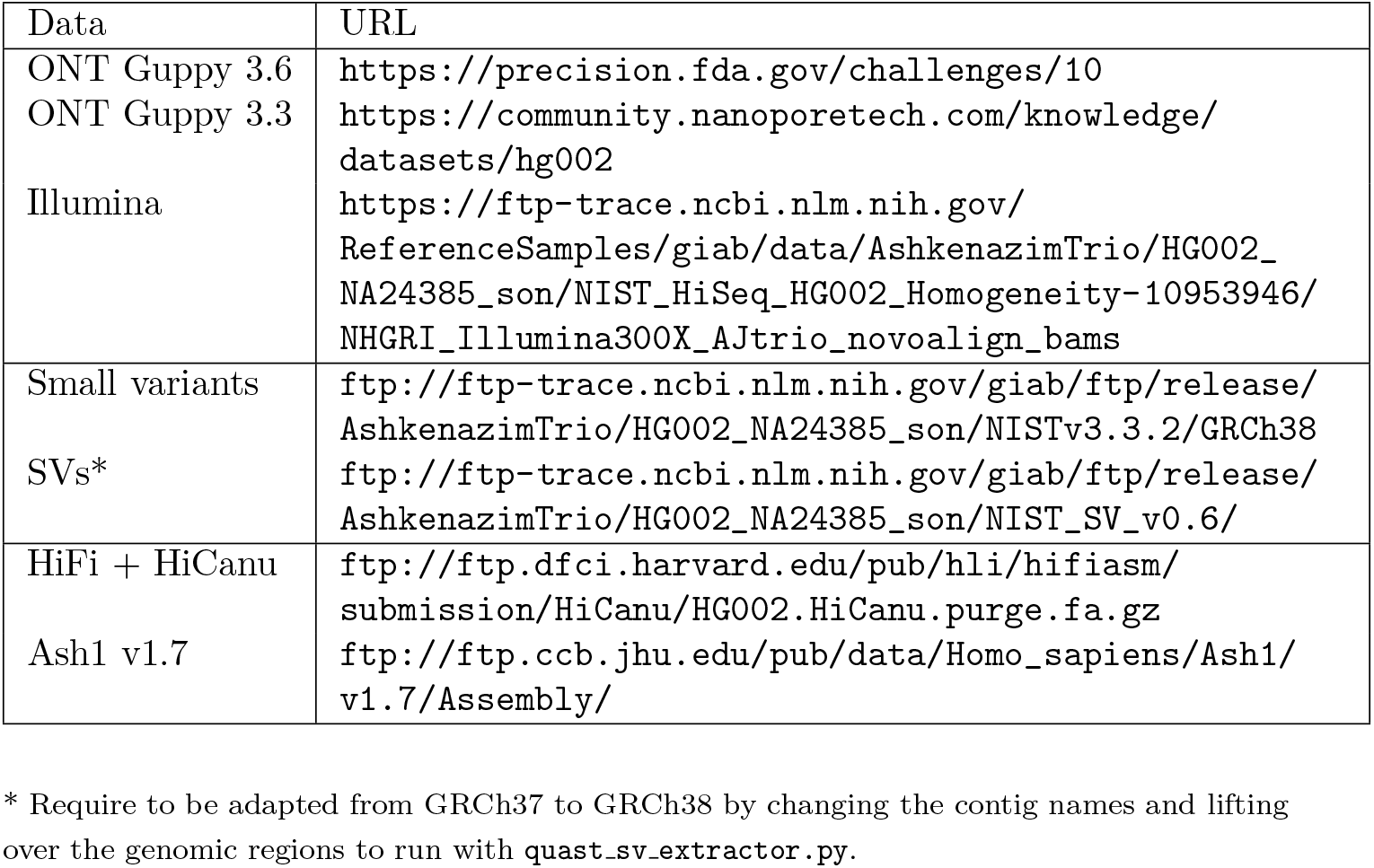

